# De novo profiling of RNA viruses in *Anopheles* malaria vector mosquitoes from forest ecological zones in Senegal and Cambodia

**DOI:** 10.1101/464719

**Authors:** Eugeni Belda, Ferdinand Nanfack Minkeu, Karin Eiglmeier, Guillaume Carissimo, Inge Holm, Mawlouth Diallo, Diawo Diallo, Amélie Vantaux, Saorin Kim, Igor V. Sharakhov, Kenneth D. Vernick

**Affiliations:** Unit of Insect Vector Genetics and Genomics, Department of Parasites and Insect Vectors, Institut Pasteur, Paris, France; CNRS Unit of Evolutionary Genomics, Modeling, and Health (UMR2000), Institut Pasteur, Paris, France; Integromics Unit, Institute of Cardiometabolism and Nutrition, Assistance Publique Hôpitaux de Paris, Pitié-Salpêtrière Hospital, Paris, France; Graduate School of Life Sciences ED515, Sorbonne Universités UPMC Paris06, 4 Place Jussieu, 75252 Paris, France.; Laboratory of Microbial Immunity, Singapore Immunology Network, Agency for Science, Technology and Research (A(∗)STAR), Singapore; Institut Pasteur de Dakar, Dakar, Senegal; Institut Pasteur of Cambodia, Phnom Penh, Cambodia; Department of Entomology, Virginia Polytechnic Institute and State University, Blacksburg VA, USA

**Author notes:** Author email: EB, FNM, KE, GC, IH, MD, DD, AV, SK, IVS, KDV.

**Keywords:** virus genome assembly, insect specific virus, RNA virus, *Anopheles*, malaria vector, virome

## Abstract

**Background:** Mosquitoes are colonized by a large but mostly uncharacterized natural virome of RNA viruses. *Anopheles* mosquitoes are efficient vectors of human malaria, and the composition and distribution of the natural RNA virome may influence the biology and immunity of *Anopheles* malaria vector populations.

**Results:** *Anopheles* vectors of human malaria were sampled in forest village sites in Senegal and Cambodia, including *Anopheles funestus, Anopheles gambiae* group sp., and *Anopheles coustani* in Senegal, and *Anopheles hyrcanus* group *sp., Anopheles maculatus* group *sp*., and *Anopheles dirus* in Cambodia. Small and long RNA sequences were depleted of mosquito host and de novo assembled to yield non-redundant contigs longer than 500 nucleotides. Analysis of the assemblies by sequence similarity to known virus families yielded 125 novel virus sequences, 39 from Senegal *Anopheles* and 86 from Cambodia. Important monophyletic virus clades in the *Bunyavirales* and *Mononegavirales* orders are found in these *Anopheles* from Africa and Asia. Small RNA size and abundance profiles were used to cluster non-host RNA assemblies that were unclassified by sequence similarity. 39 unclassified non-redundant contigs >500 nucleotides strongly matched a pattern of classic RNAi processing of viral replication intermediates, and 1566 unclassified contigs strongly matched a pattern consistent with piRNAs. Analysis of piRNA expression in *Anopheles coluzzii* after infection with O’nyong nyong virus (family *Togaviridae*) suggests that virus infection can specifically alter abundance of some piRNAs.

**Conclusions:** RNA viruses ubiquitously colonize Anopheles vectors of human malaria worldwide. At least some members of the mosquito virome are monophyletic with other arthropod viruses. However, high levels of collinearity and similarity of Anopheles viruses at the peptide level is not necessarily matched by similarity at the nucleotide level, indicating that *Anopheles* from Africa and Asia are colonized by closely related but clearly diverged virome members. The interplay between small RNA pathways and the virome may represent an important part of the homeostatic mechanism maintaining virome members in a commensal or nonpathogenic state, and host-virome interactions could influence variation in malaria vector competence.

## Introduction

*Anopheles* mosquitoes are the only vectors of human malaria, which kills at least 400,000 persons and causes 200 million cases per year, with the greatest impact concentrated in sub-Saharan Africa and South-East Asia [1]. In addition to malaria, *Anopheles* mosquitoes also transmit the alphavirus O’nyong nyong (ONNV, family *Togaviridae*), which is the only arbovirus known to employ *Anopheles* mosquitoes as the primary vector [2, 3].

*Anopheles* mosquitoes harbor a diverse natural virome of RNA viruses [4-7]. A recent survey found evidence of at least 51 viruses naturally associated with *Anopheles* [2]. The *Anopheles* virome is composed mainly of insect specific viruses (ISVs) that multiply only in insects, but also includes relatives of arboviruses that can replicate in both insects and vertebrate cells.

Culicine mosquitoes in the genera *Aedes* and *Culex* transmit multiple arboviruses such as dengue (DENV, family *Flaviviridae*) Zika (ZIKV, family *Flaviviridae*), chikungunya (CHIKV, family *Togaviridae*) and others, but do not transmit human malaria. This apparent division of labor between culicine and *Anopheles* mosquitoes for transmission of arboviruses and Plasmodium, respectively, has led to a relative lack of study about *Anopheles* viruses. *Anopheles* viruses have been discovered by isolation from cultured cells exposed to mosquito extract, serology, specific amplification and sequencing, and more recently, deep sequencing and de novo assembly [2]. Although this work has increased the number of ISVs discovered in Anopheles, it appears that there are many still unknown.

Here, we assembled small and long RNA sequences from wild *Anopheles* mosquitoes captured in forest ecologies in central and northern Cambodia and eastern Senegal. The sites are considered disease emergence zones, with high levels of fevers and encephalopathies of unknown origin. Sequence contig evidence of a number of novel RNA viruses and variants was detected, and potentially many unclassified viruses.

It is likely that persistent exposure to ISVs, rather than the relatively infrequent exposure to arboviruses such as ONNV, has been the main evolutionary pressure shaping *Anopheles* antiviral immunity. *Anopheles* resistance mechanisms against arbovirus infection may be quite efficient, based on their lack of virus transmission despite highly anthropophilic feeding behavior, including on viremic hosts. Nevertheless, ONNV transmission is the exception that indicates arbovirus transmission by *Anopheles* is possible, so it is a biological puzzle that transmission is apparently restricted to just one virus. Identifying the complement of natural viruses inhabiting the *Anopheles* niche will help clarify the biology underlying the apparent inefficiency of arbovirus transmission by *Anopheles*, and may suggest new tools to raise the barrier to arbovirus transmission by the more efficient *Aedes* and *Culex* vectors.

## Results

### Mosquito species estimation

Metagenomic sequencing of long and small fractions of RNA was carried out for four biological replicates pools of mosquitoes from Ratanakiri and Kampong Chnang provinces in central and northern Cambodia near the border with Laos, and four replicate pools from Kedougou in eastern Senegal near the border with the Republic of Guinea (Conakry). Mosquito species composition of sample pools was estimated using sequences of transcripts from the mitochondrial cytochrome c oxidase subunit 1 (COI) gene, which were compared with *Anopheles* sequences from the Barcode of Life COI-5P database (Figure 1, Additional File 1: Table S1). In the Senegal samples, the most frequent mosquito species were *Anopheles rufipes*, *Anopheles funestus, Anopheles gambiae* group sp., and *Anopheles coustani*, which are all human malaria vectors, including the recently incriminated *An. rufipes* [8]. In the Cambodia samples, the most frequent species were *Anopheles hyrcanus* group *sp., Anopheles maculatus* group *sp*., *Anopheles karwari, Anopheles jeyporeisis, Anopheles aconitus* and *Anopheles dirus*. All are considered human malaria vectors [9-12]. Elevated rates of human blood-feeding by a mosquito species is a prerequisite for malaria vectorial capacity [13], and therefore the main *Anopheles* species sampled for virome discovery in this study display consistently high levels of human contact in nature.

**Figure 1.**
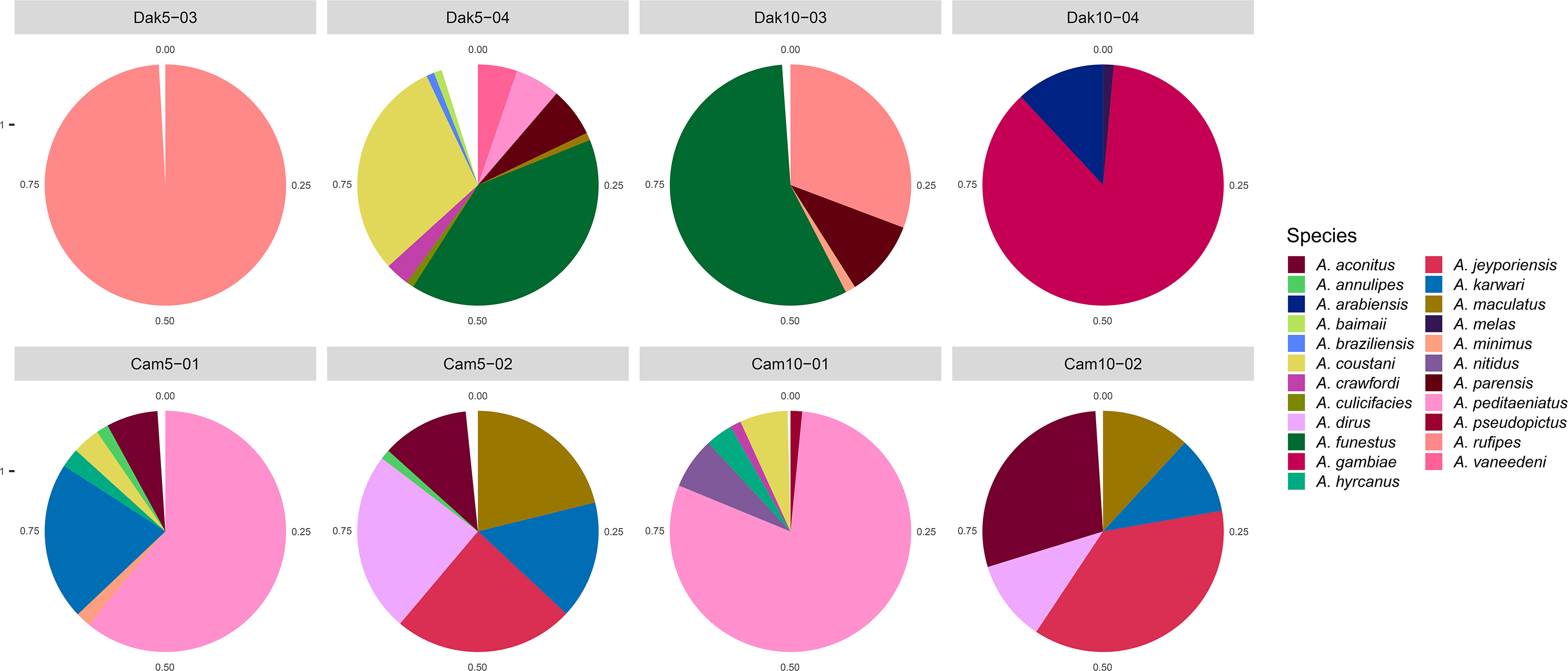
Taxonomic profile of *Anopheles* sample pools. Relative abundances of *Anopheles* species were computed from mapping of long-RNA reads over mitochondrial cytochrome C oxidase subunit I gene sequences (COI-5P) from Barcode of Life Database. Taxa represented by >100 sequence reads and 1% frequency in the sample pool were plotted in pie charts. White wedges represent the proportion of sequence matches present at less than 1% frequency. All data are presented in tabular form in Additional File: Table S1.

### Virus discovery by de novo RNAseq assembly and classification by sequence similarity

Small and long RNA reads were de novo assembled after removal of mosquito sequences. Non-redundant contigs longer than 500 nucleotides from assemblies of both countries, Cambodia and Senegal, were used to search the GenBank protein sequence database using BLASTX with an e-value threshold of 1e-10. This allowed identification of 125 novel assembled virus sequences, 39 from the Senegal samples (virus ID suffix “Dak”, Table 1), and 86 from the Cambodia samples (virus ID suffix “Camb”, Table 2), possibly pointing to higher viral diversity in mosquitoes from Cambodia. Some of the 125 virus sequences showed remote similarity by BLASTX to 24 reference viruses in GenBank that include ssRNA-negative strand viruses of the families *Orthomyxoviridae*, *Rhabdoviridae* and *Bunyaviridae*, ssRNA positive-strand viruses of the families *Virgaviridae*, *Flaviviridae* and *Bromoviridae*, dsRNA viruses of the family *Reoviridae* and multiple unclassified viruses of both ssRNA and dsRNA types (Table 3). Most of these remote similarities were with viruses characterized in a recent virus survey of 70 different arthropod species collected in China [14], which emphasizes the importance of high throughput surveys of arthropod virosphere in the identification of viruses associated with different arthropod species.

**Table 1.**
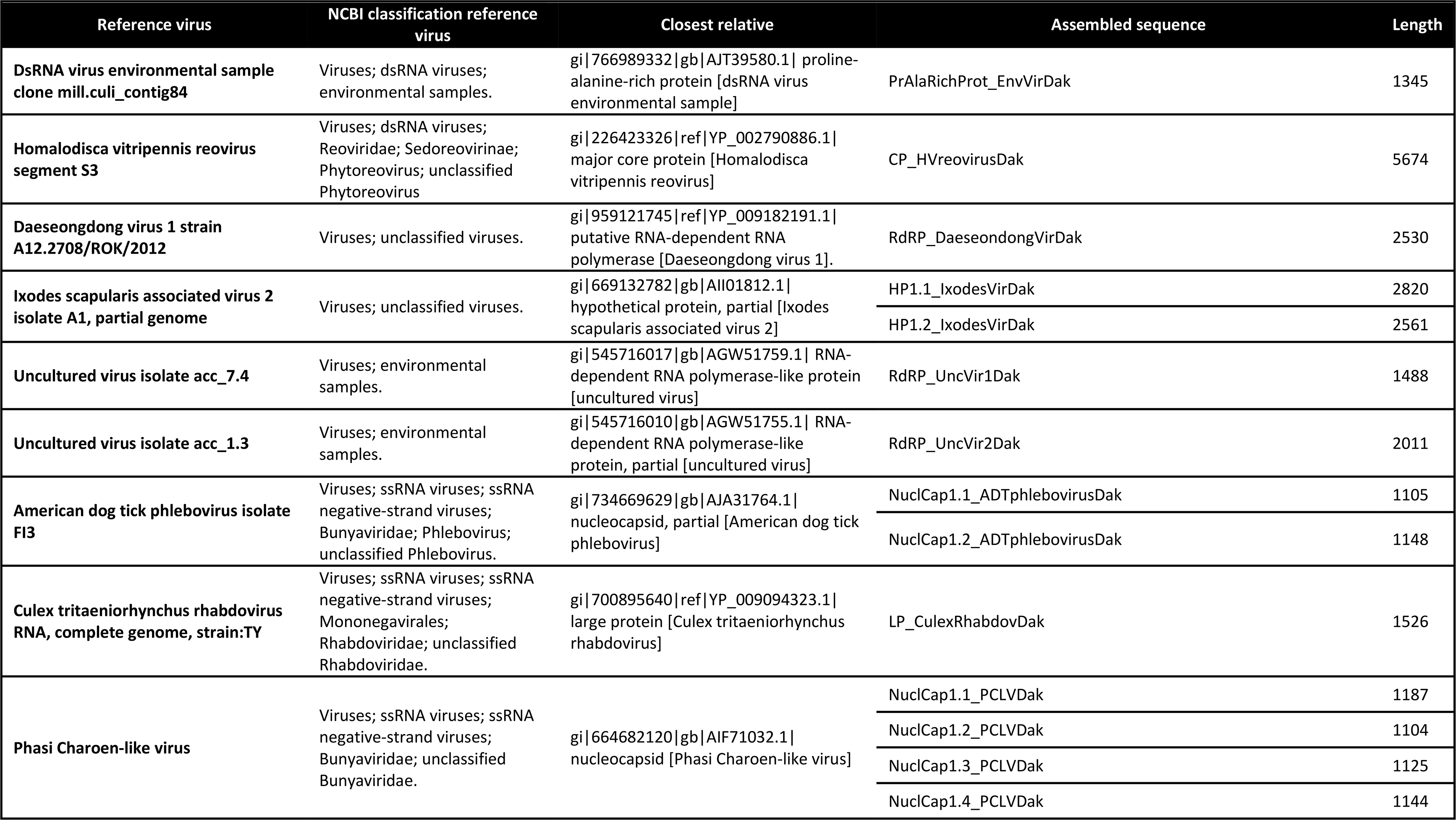
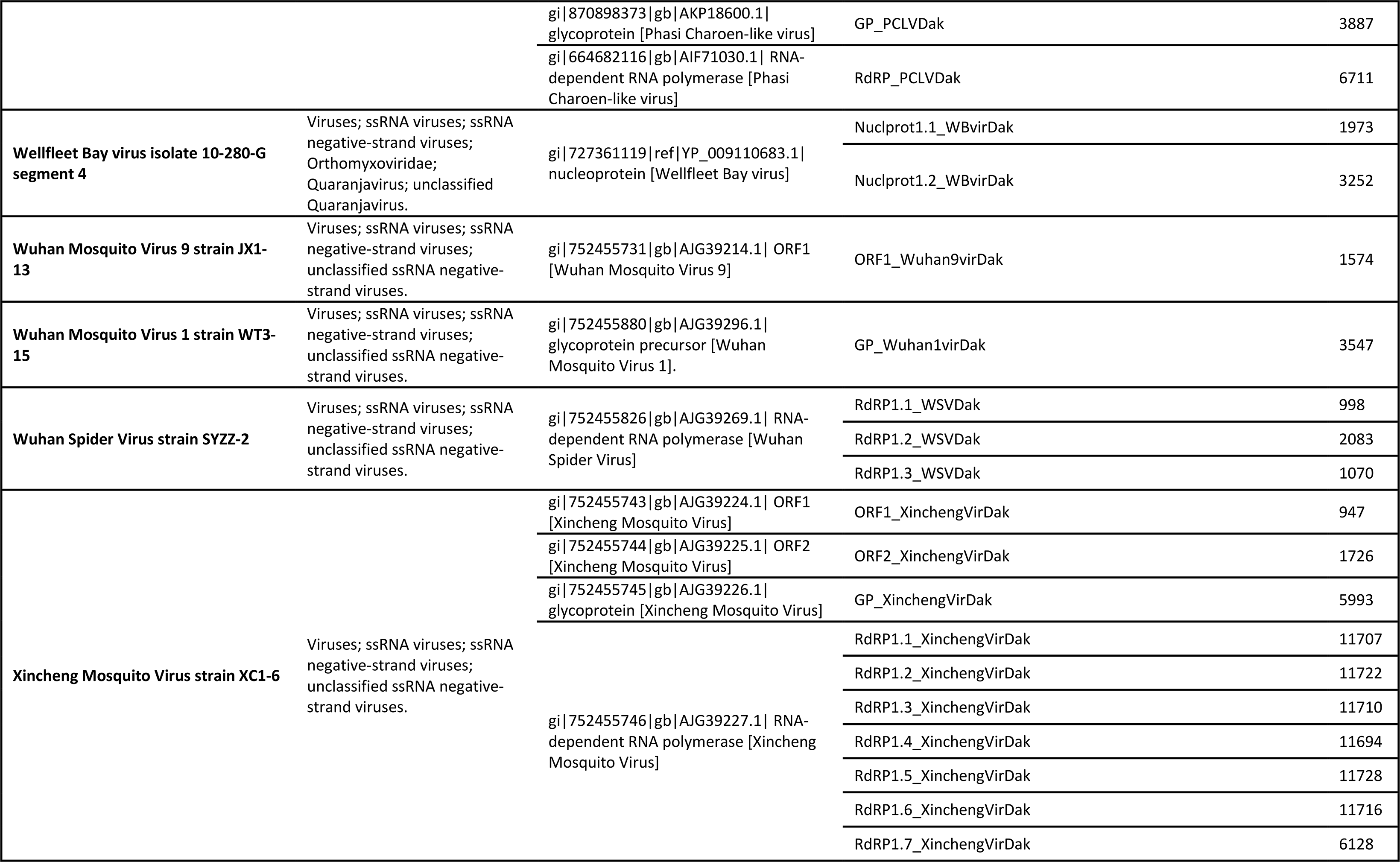
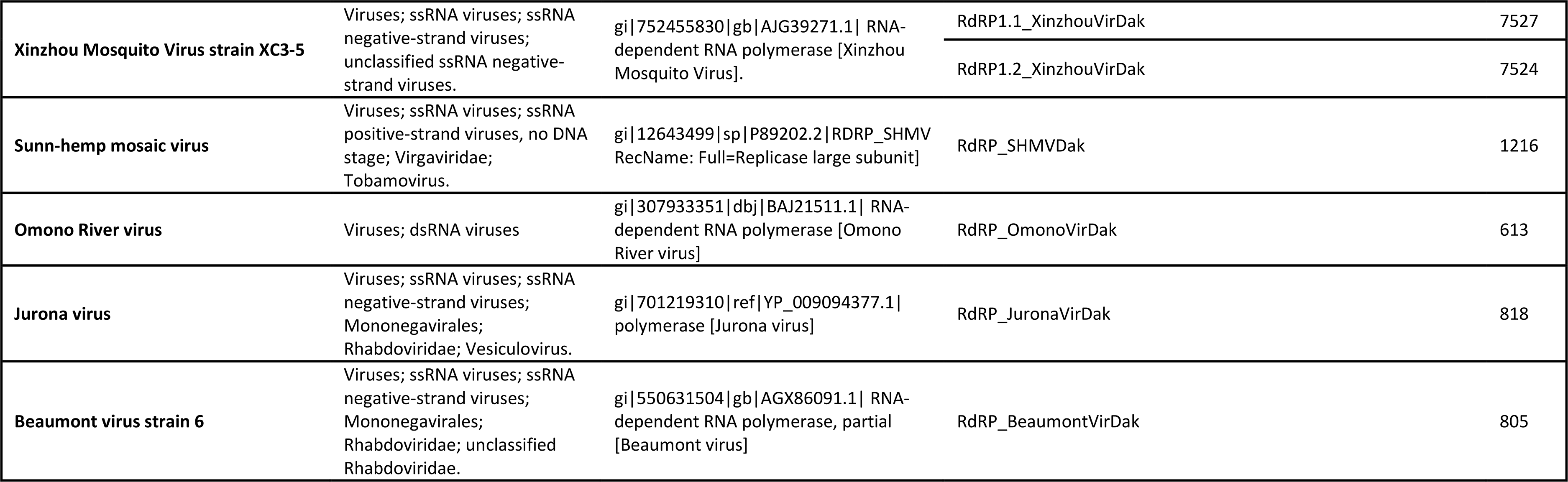
Summary of virus assemblies, Senegal *Anopheles* sample pools.

**Table 2.**
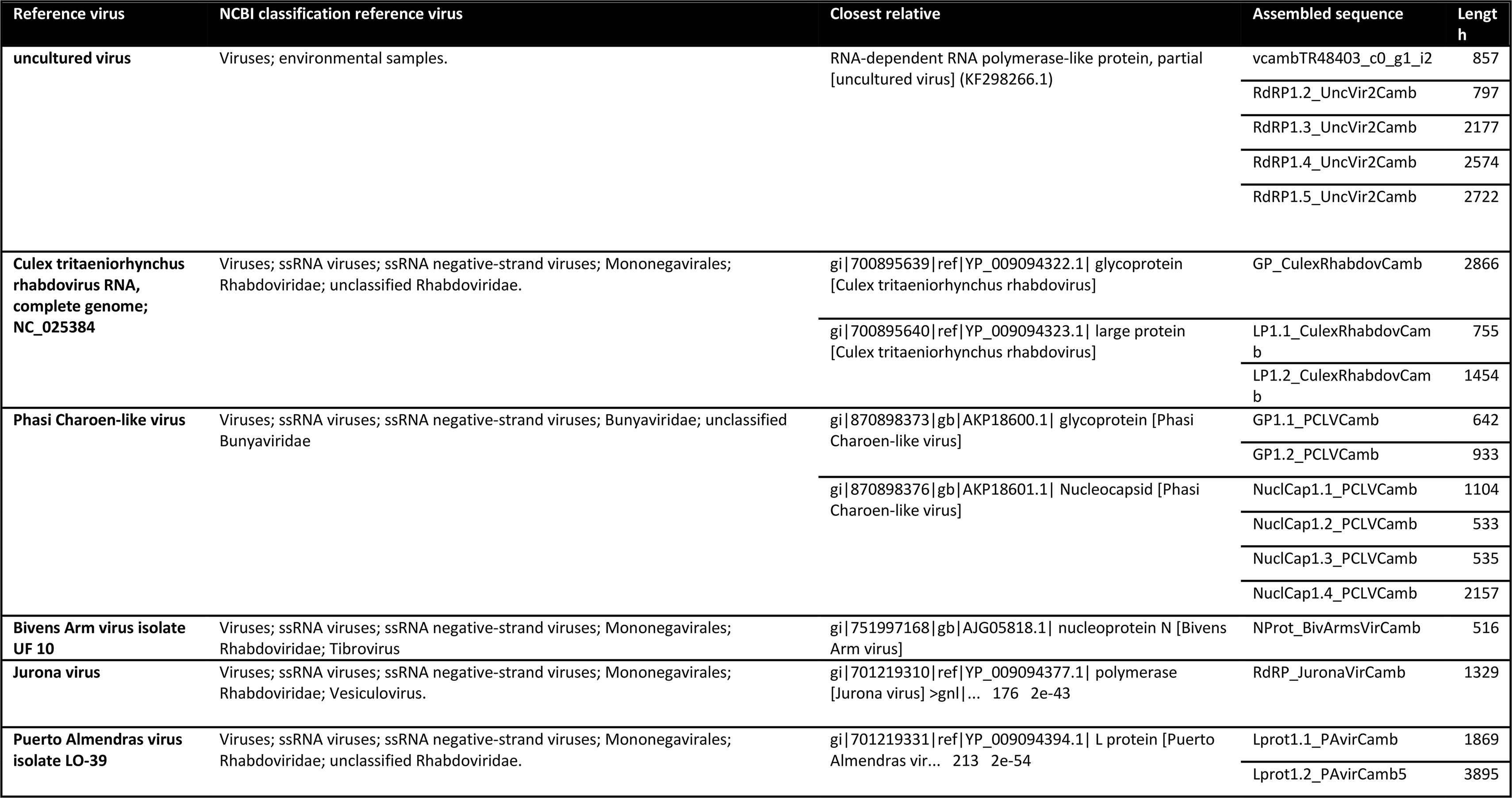
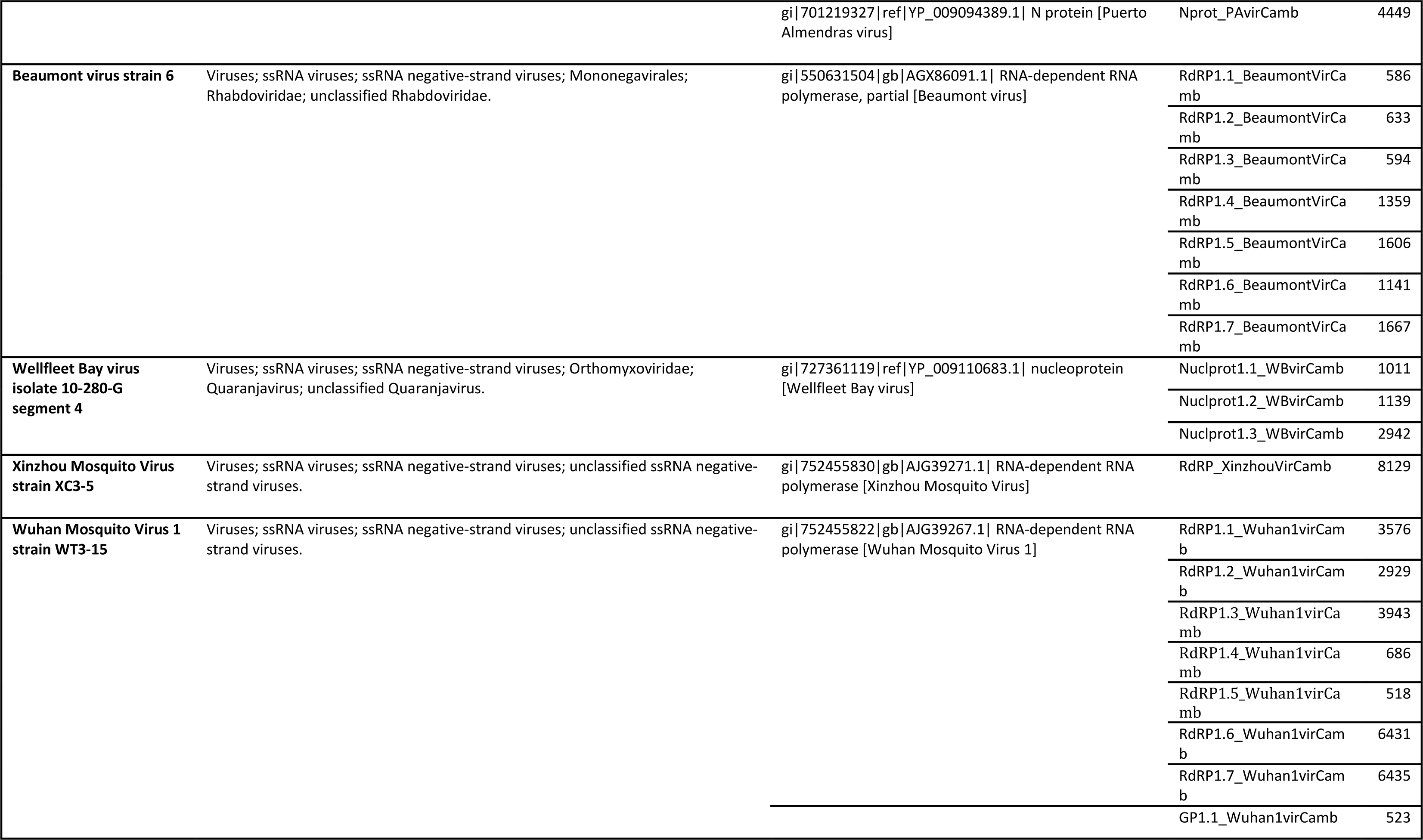
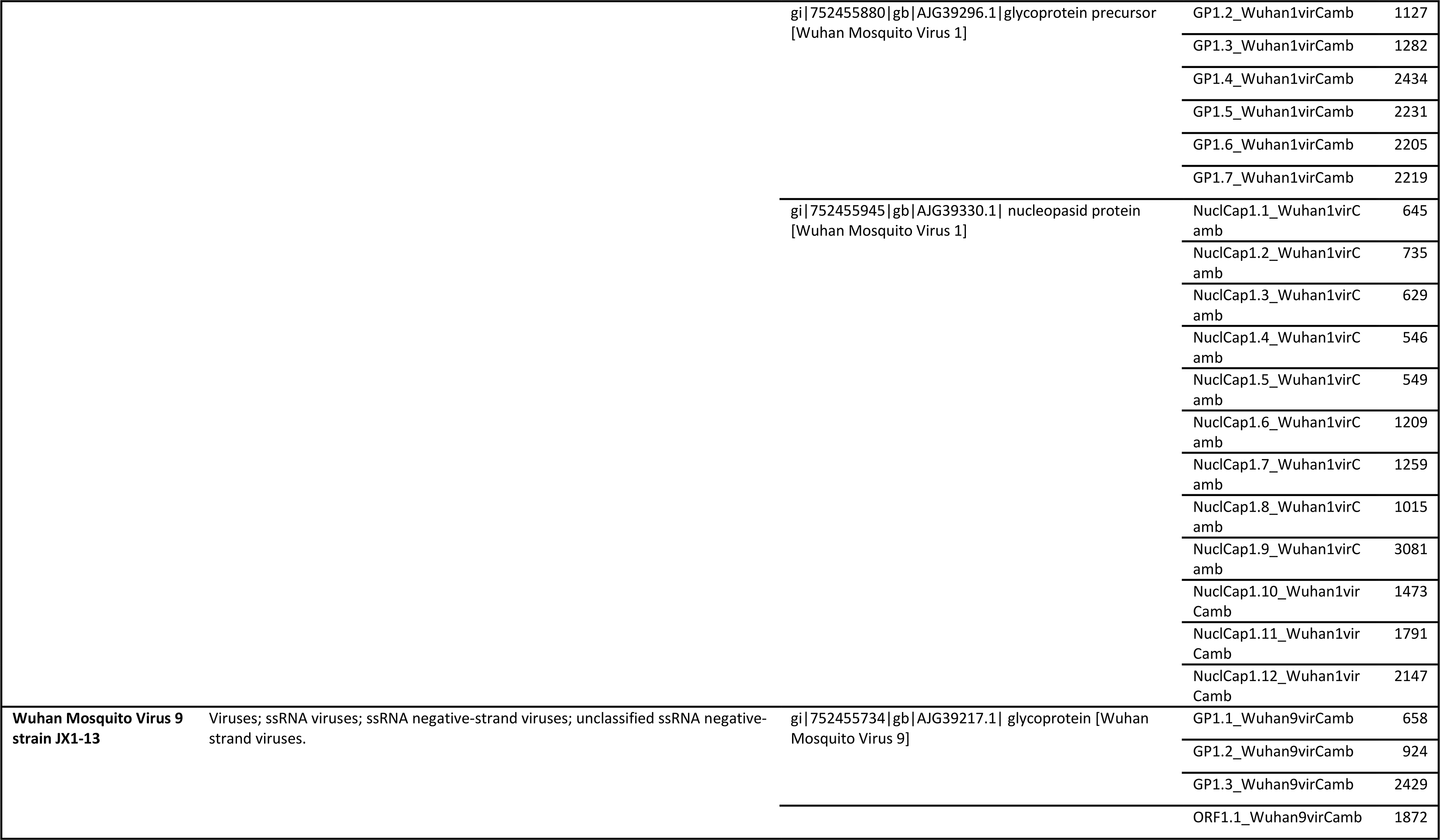
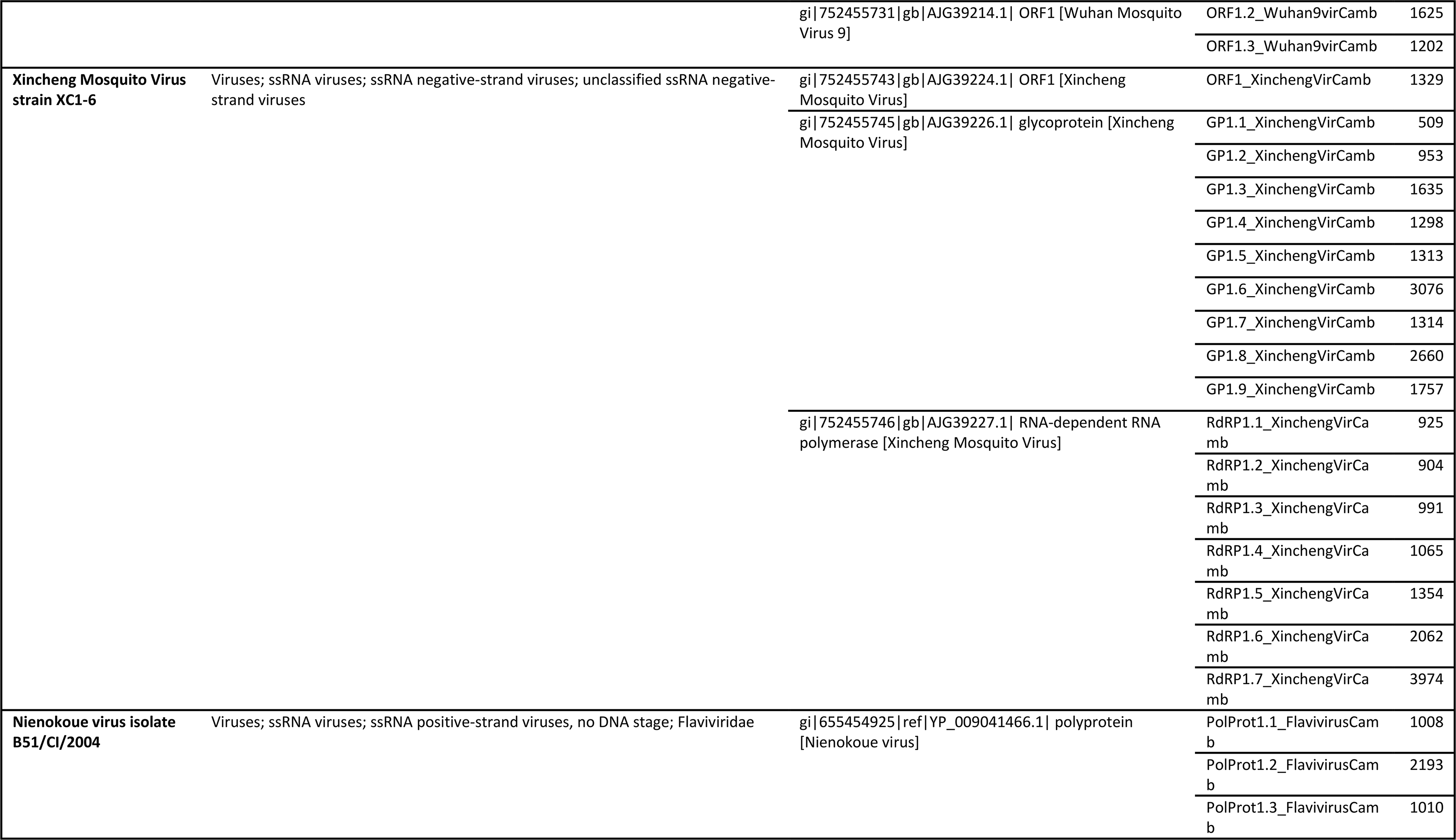
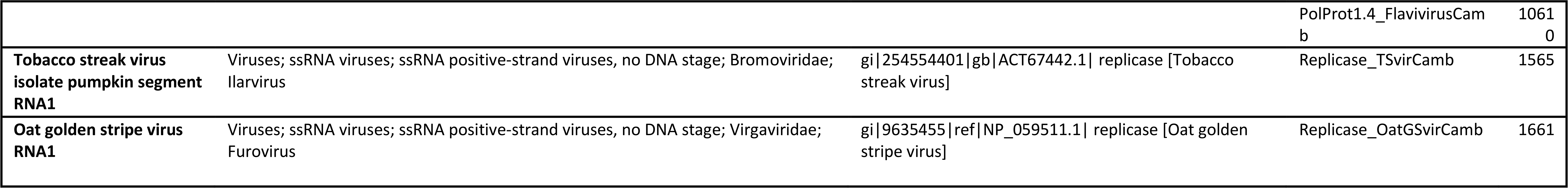
Summary of virus assemblies, Cambodia *Anopheles* sample pools.

**Table 3.**
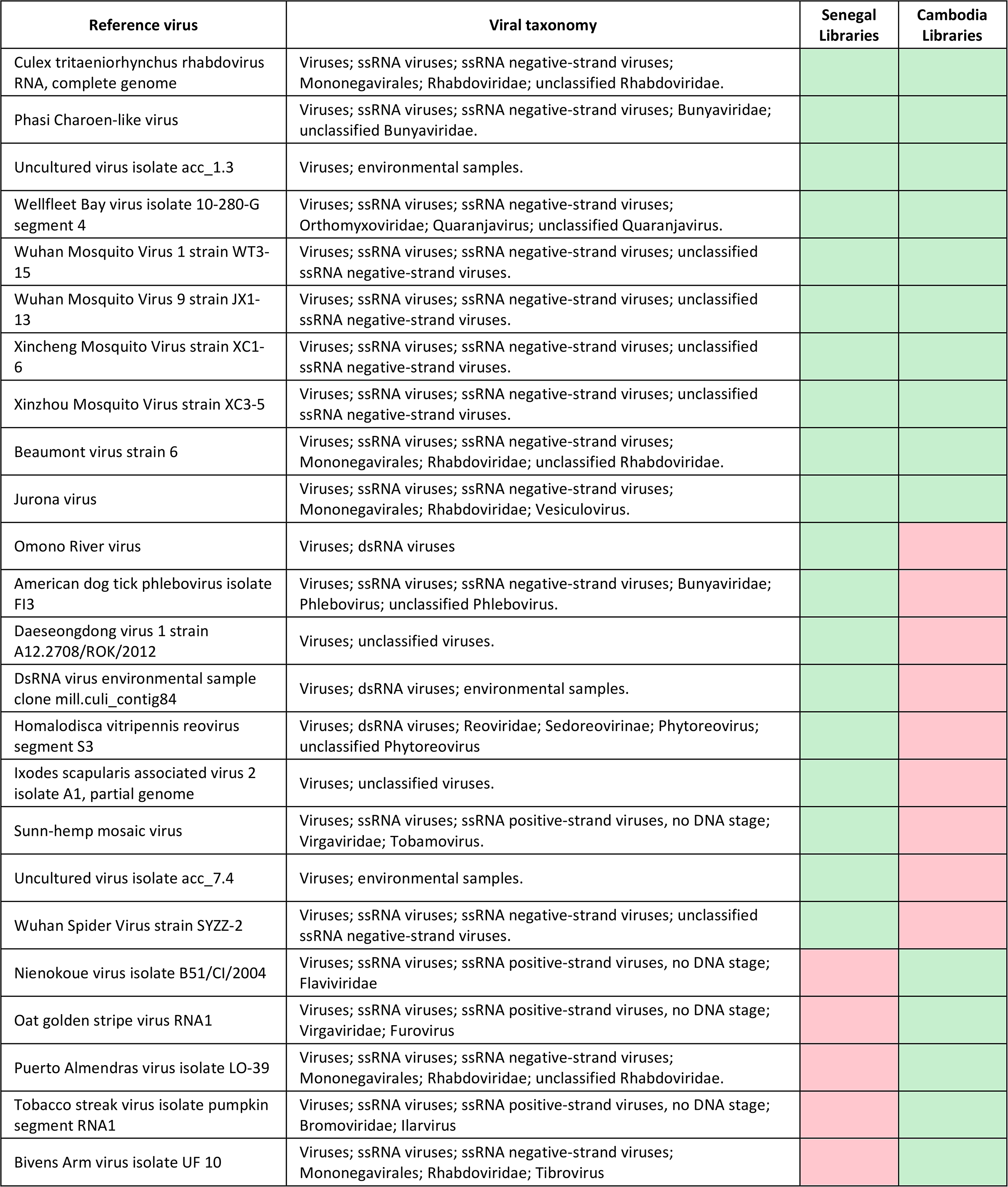
Similarity of Senegal and Cambodia virus assemblies by BLASTX to 24 reference viruses in GenBank. 10 targets are shared, 9 are Senegal-specific, and 5 are Cambodia-specific.

In order to place these 125 novel virus assemblies in an evolutionary context, phylogenetic trees were constructed from conserved regions of the RNA-dependent RNA polymerase gene annotated in the 125 virus sequences, along with related virus sequences from GenBank. This allowed the placement of 44 of the 125 assembled viruses in phylogenetic trees, revealing clusters of highly related viruses in the analyzed wild *Anopheles*. Notable examples include five novel virus assemblies from Cambodian *Anopheles* placed near Wuhan Mosquito Virus 1 in a monophyletic group of the Phasmavirus clade (*Bunyavirales*) (Figure 2). Also, within the order *Mononegavirales*, 14 novel *Anopheles* virus assemblies (7 from Cambodia and 7 from Senegal) formed a monophyletic group that includes Xincheng Mosquito Virus and Shungao Fly Virus. Finally, 10 novel virus assemblies (9 from Cambodia, 1 from Senegal) formed a monophyletic group that includes Beaumont Virus and a rhabdovirus from *Culex tritaeniorhynchus* within the Dimarhabdovirus clade (Figure 3A). TBLASTX comparisons of virus sequences in these groups with the closest reference viruses in the phylogenetic trees showed high levels of collinearity and similarity at protein level that was not matched by comparable levels of similarity at the nucleotide level, indicating that populations of closely related but diverged viruses colonize *Anopheles* from widely separated geographic locations (Figure 3B).

**Figure 2.**
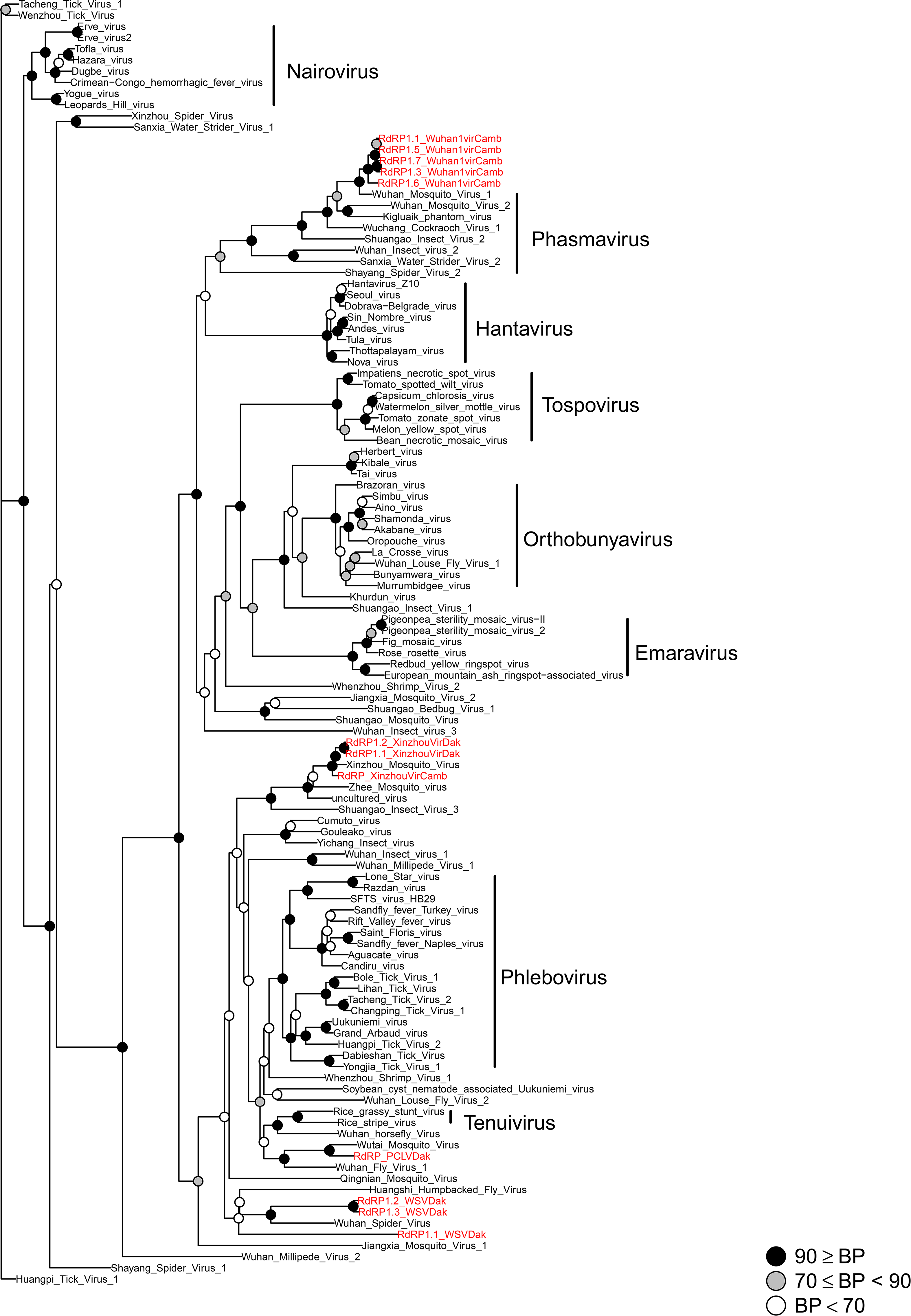
Phylogenetic tree of reference and novel virus assemblies from the *Bunyaviridae* family. Novel viruses characterized from Cambodia and Senegal*Anopheles* sample pools (red labels) are placed within the Phasmavirus clade and in a basal position of the Phebovirus-Tenuivirus clade.

**Figure 3.**
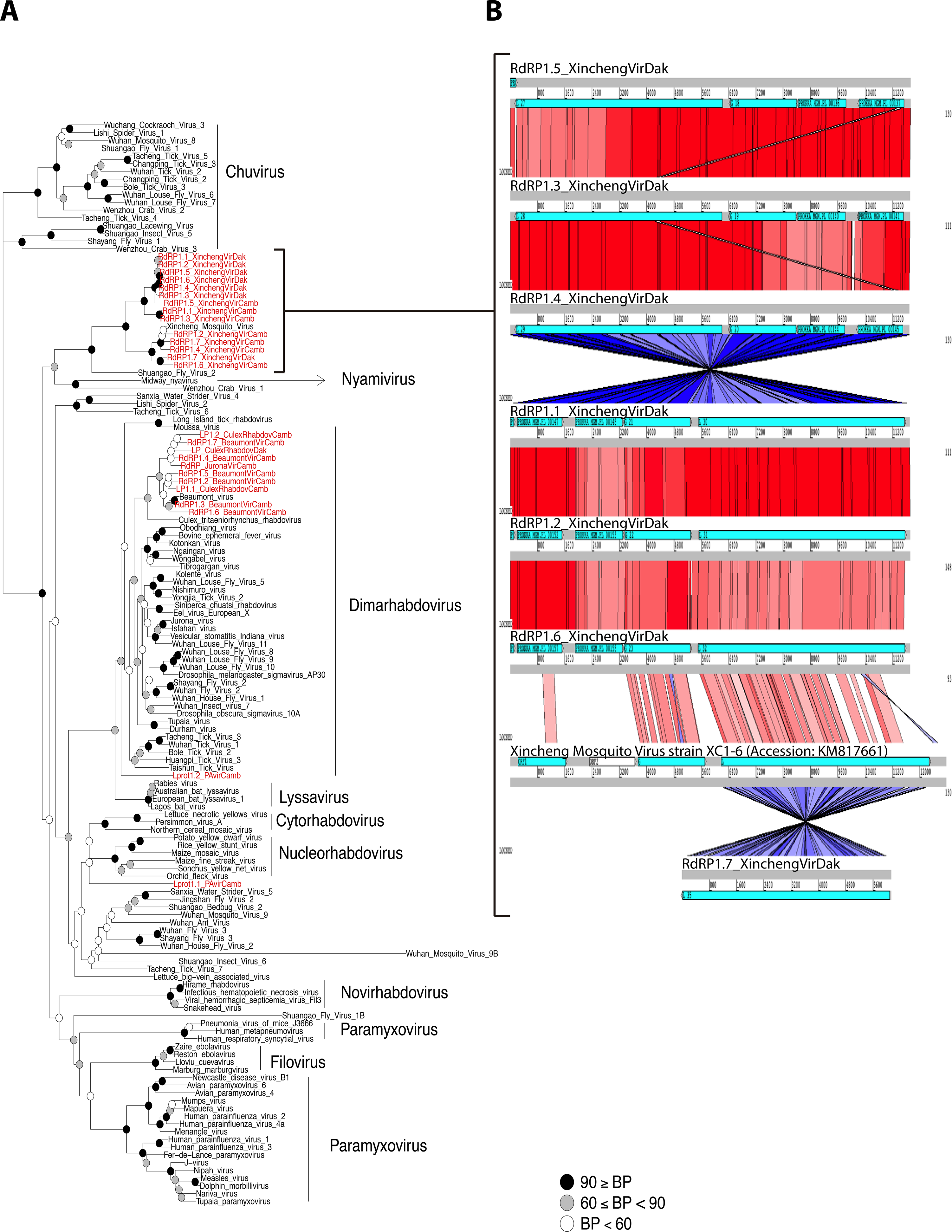
Phylogenetic tree of reference and novel virus assemblies from the *Mononegavirales* order. **A)**Novel virus assemblies characterized from Cambodia and Senegal *Anopheles* sample pools (red labels) are predominantly placed within the Dimarhabdovirus clade and as close relative of the Nyamivirus clade. **B)** In this latter group close to Nyamivirus, the novel virus assemblies identified are close relatives of Xincheng mosquito virus, sharing a high degree of genome collinearity based on TBLASTX comparisons of novel and reference Xinxeng mosquito reference sequences.

### Quantification of novel virus sequences in mosquito sample pools

In order to evaluate the prevalence of novel virus sequences across the analyzed mosquito samples, host-filtered small and long RNA reads were mapped over the 125 novel virus sequences identified by de novo sequence assembly. Based on long RNAseq reads, the abundance profiles of the 125 virus assemblies display a non-overlapping distribution across different sample pools, and virus sequences can be localized to particular sample pools from the abundance profiles (Figure 4, left panel). This probably indicates a patchy prevalence and abundance of the different viruses among individual mosquitoes, such that an individual mosquito highly infected with a given virus could potentially generate a strong signal for the virus in the sample pool. The sample pools from Cambodia share a higher fraction of common viruses, while there is less overlap in virus abundance distribution across sample pools from Senegal. The representation of virus distribution based on small RNA sequence reads displayed profiles broadly similar to the long RNA-based abundance distribution (Figure 4, right panel). This observation may be consistent with the expectation that small RNA representation is a signature of virus double-stranded RNA (dsRNA) processing by the mosquito RNA interference (RNAi) machinery [15], and therefore was specifically examined next.

**Figure 4.**
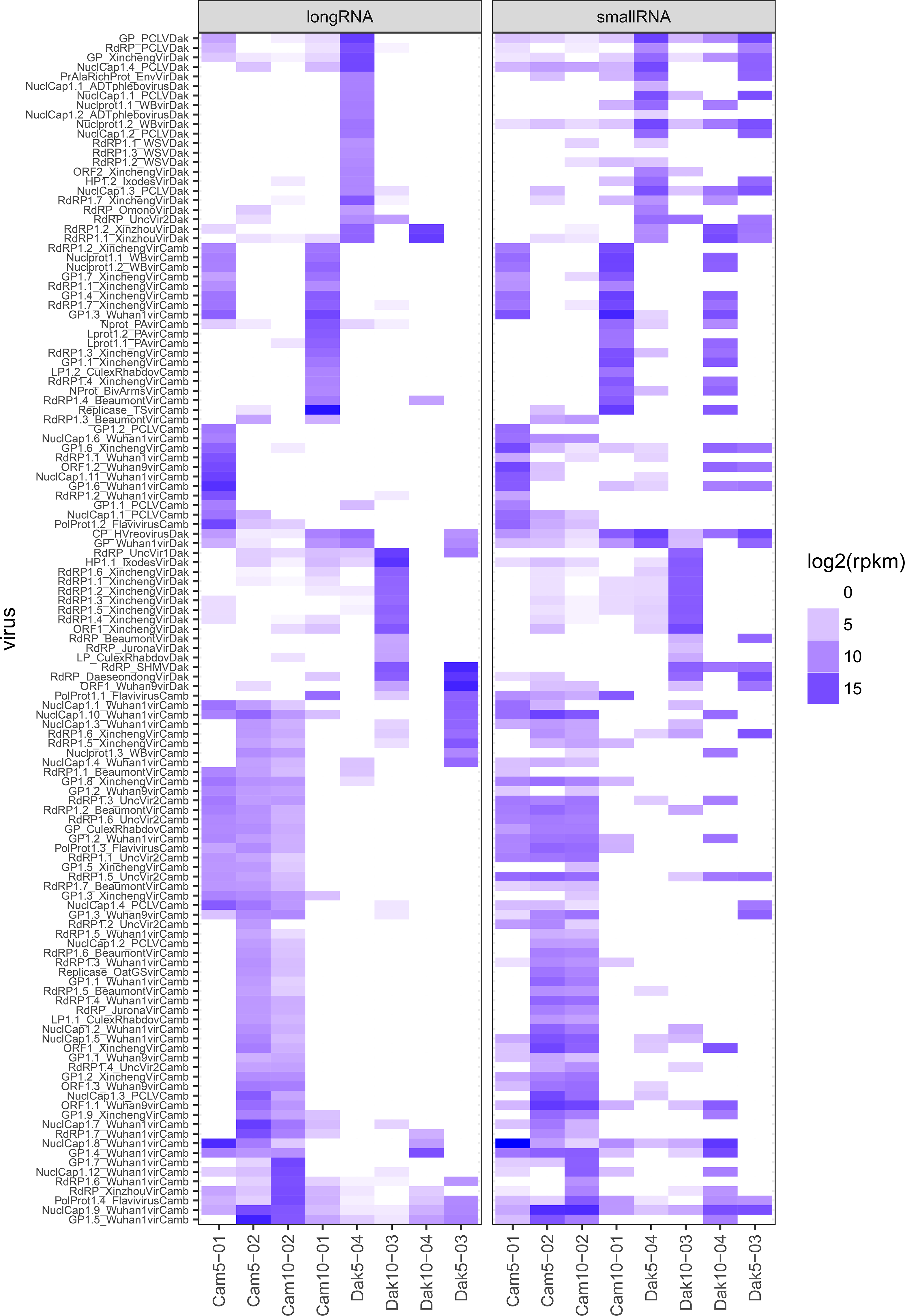
Viral abundance profiles across mosquito sample pools based on small and long RNA sequence mapping. Heatmap of log2-transformed reads per kilobase per million reads (RPKM) abundance values of novel virus assemblies identified from Cambodia and Senegal pools based on long and small RNA sequence libraries. Broadly similar viral abundance profiles are observed for different pools based on small and long RNA sequence data. Representation of particular viruses is uneven among pools, possibly indicating inter-individual mosquito differences for virus carriage.

### Small RNA size profiling

The processing of virus sequences by small RNA pathways of the insect host generates diagnostic patterns of small RNA read sizes from different viruses. In order to evaluate this phenomenon in the 125 novel virus assemblies characterized by sequence similarity in the analyzed sample pools, small RNA reads that mapped to each virus assembly were extracted, and their size distributions were normalized with a z-score transformation. This allowed comparison of the z-score profiles among virus assemblies by pairwise correlation analysis and hierarchical clustering. The relationship between the small RNA profiles of the different viruses could then be visualized as a heat map. The results of this analysis revealed the presence of four major groups of virus sequences based on small RNA size profiles (Figure 5). Cluster 1 consists of 7 virus assemblies generating small RNAs predominantly in the size range of 23-29 nt mapping over the positive, and to a lesser extent negative, strand. Cluster 2 includes 7 viruses, all from Senegal, and displays a similar size profile as viruses of Cluster 1 with reads in the 23-29 nt size range, but also with a higher frequency of 21 nt reads mapping over the positive and negative strands, emblematic of virus cleavage through the mosquito host RNAi pathway. Cluster 3 includes 15 viruses that exhibit the classic pattern of viruses processed by the host RNAi pathway, with predominantly reads of 21 nt in length mapping over virus positive and negative strands (Additional File 2: Figure S1). Finally, Cluster 4 includes 59 viruses with small RNA size profiles dominated by reads of 23-29 nt mapping predominantly over the negative strand of virus sequences. Because of the strong strand bias of small RNAs observed, this pattern could correspond to degradation products of virus RNAs, although alternatively, there appears to be size enrichment in the 27-28 nt size peaks characteristic of PIWI-interacting RNAs (piRNAs).

**Figure 5.**
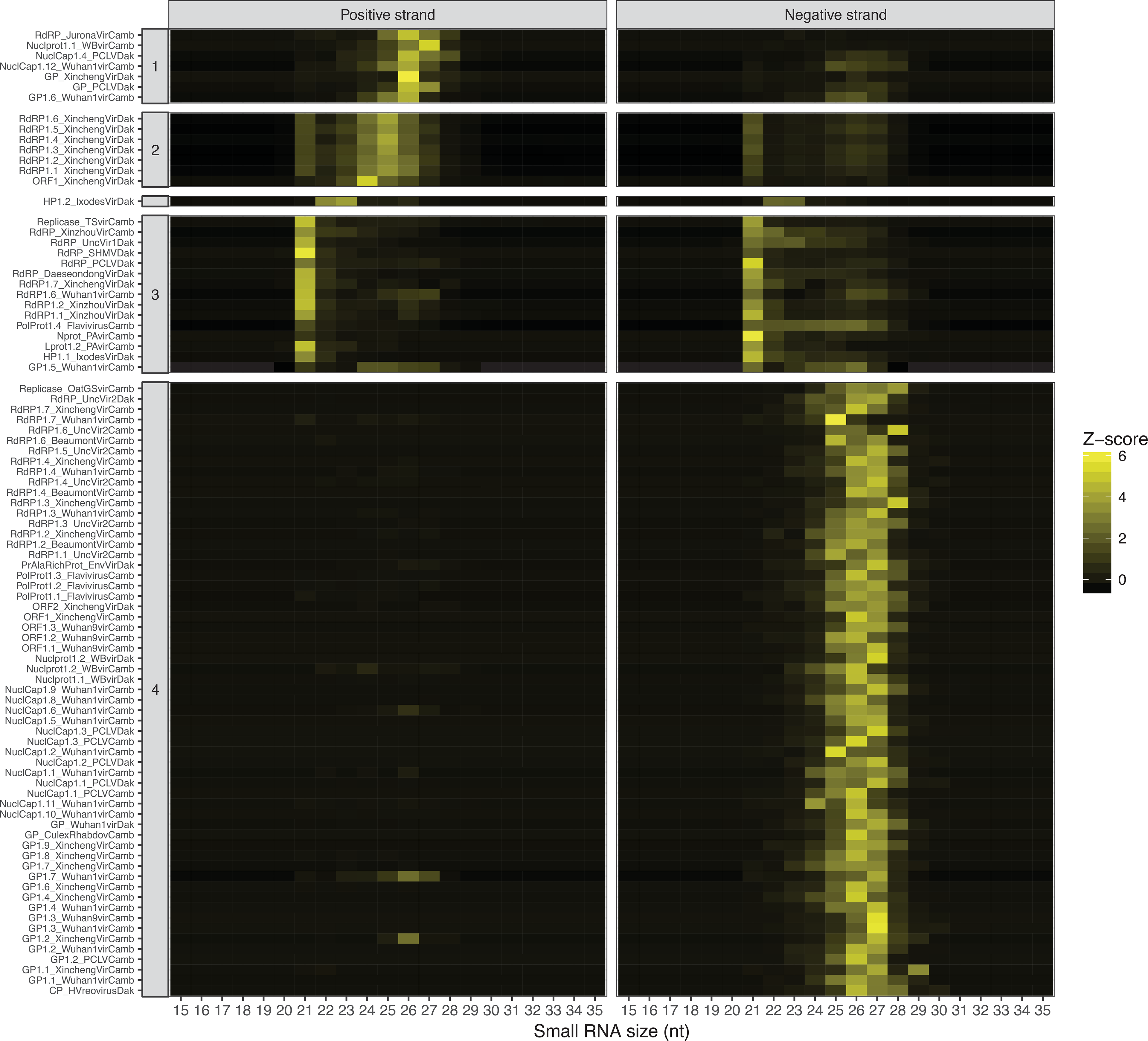
Small RNA size profiles of novel virus assemblies from Cambodia and Senegal sample pools. Hierarchical clustering of novel virus assemblies based on Pearson correlation of z-score transformed small RNA size profiles (the frequency of small RNA reads of size 15 to 35 nucleotides that maps over the positive and negative strand of the reference sequence). Four main clusters were defined based on these small RNA size profiles, among which the classical siRNA size profile (21 nt reads mapping over positive and negative strand) is represented in the Cluster 3.

### Viral origin of unclassified transcripts by small RNA size profiling

A major drawback of sequence similarity-based identification of novel viruses in de novo sequence assemblies is the dependence of detection upon existing records of close relatives in public databases. It was proposed that the small RNA size profiles of arthropod-derived viruses detected by sequence similarity could be used as signature to recruit unclassified contigs from de novo sequence assemblies of potential viral origin [15]. We implemented this strategy in order to identify additional sequences of putative viral origin in the set of 2114 contigs with at least 100 small RNA sequence reads left unclassified by sequence similarity searching.

Of these unclassified contigs, a likely viral origin is supported for 4 and 35 contigs that display strong association by small RNA profile with Cluster 2 and Cluster 3, respectively (Spearman correlation>0.9, Additional File 3: Figure S2). These clusters display small RNA size profiles mapping to both genome strands, and characteristic of classic RNAi processing of viral dsRNA replication intermediates. Thus, in addition to the 125 novel virus assemblies classified by sequence similarity to known viruses, 39 unclassified novel *Anopheles* virus assemblies were identified, without sequence similarity to identified viruses but meeting the quality criteria of non-redundant assemblies longer than 500 nucleotides. Further work will be necessary to characterize the biology of these unclassified novel virus assemblies.

Of the other assemblies unclassified by sequence similarity, 1566 showed strong associations between their small RNA size profiles and the small RNA size profiles of virus contigs detected by sequence similarity (Spearman correlation>0.9). Among these, the majority were associated with Cluster 4 virus assemblies (1219 unclassified contigs) and to less extent with Cluster 1 (309 unclassified contigs). Both clusters were characterized by a strong bias towards reads from a single strand (positive for Cluster 1 and negative for Cluster 4).

To evaluate how specific these latter profiles of 1219 and 309 contigs are for virus-related sequences, we designed a reconstruction control experiment using the same small RNA size profiling and clustering analysis as above, but instead using 669 RNA contigs known to map to the mosquito reference assembly, thus strictly of host origin. As above, contigs with at least 100 small RNA sequence reads were used. 561 of these mosquito contigs could be grouped with small RNA size profiles of virus contigs (Spearman correlation>0.9), most of them (98.21%) with Cluster 4 (78.6%) and Cluster 1 (19.6%) profiles.

However, many somatic piRNAs map to only one strand in Drosophila and other arthropods [16, 17]. Notably, many virus-related piRNAs in *Aedes*, which are largely ISV-derived, mainly map only to the virus strand antisense to the viral ORF [18]. In *An. coluzzii*, about half of expressed piRNAs display a strong or exclusive strand bias, which is a greater proportion of unidirectional piRNAs than *Drosophila* [19]. Until the current study, *Anopheles* piRNAs have not previously been examined for relatedness to ISVs. Overall, these results are probably most consistent with an interpretation that RNA profile Cluster 1 and Cluster 4 detect strand-biased piRNAs derived from the natural ISV virome of wild *Anopheles*. On that interpretation, the above host-sequence control contigs that share the Cluster 1 and Cluster 4 RNA profiles are most likely also piRNAs, but instead derived from endogenous host templates. Previous results showed that most *An. coluzzii* piRNAs target long-terminal repeat retrotransposons and DNA transposable elements [19]. Our current results add wild ISVs as a likely source of template for *Anopheles* piRNA production, and indicate that further work is warranted in the interpretation of small RNA profiles for discovery of unclassified viruses. Our results also suggest the possibility that piRNAs may be involved in *Anopheles* response to viruses, a phenomenon found for only *Aedes* among a wide range of arthropods, but *Anopheles* were not yet tested [17].

### O’nyong nyong alphavirus infection influences expression of piRNAs in *Anopheles coluzzii*

piRNAs are endogenous small noncoding RNAs of about 24-30 nt that ensure genome stability by protecting it from invasive transposable elements such as retrotransposons and repetitive or selfish sequences [17]. In addition, in *Aedes* mosquito cells, piRNAs can probably mediate responses to arboviruses or ISVs [17, 18, 20, 21]. *Anopheles* mosquitoes express piRNAs from genomic piRNA clusters [19, 22], but piRNA involvement in response or protection to virus infection in *Anopheles* has not been reported to our knowledge. To examine the potential that *Anopheles* piRNAs could be involved in response to viruses, we challenged *An. coluzzii* mosquitoes with the alphavirus, ONNV by feeding an infectious bloodmeal, and sequenced small RNAs expressed during the primary infection at 3 d post-bloodmeal. Mosquitoes fed a normal bloodmeal were used as the control condition.

Analysis of the small RNA expression data using Cuffdiff and DESeq2 detected 86 potential significantly differentially expressed transcripts between ONNV infected mosquitoes and normal bloodmeal controls (Additional File 4: Table S2). Filtering for appropriate length of contiguous expressed region for piRNA <40 nt, and high abundance of expression in ONNV and control samples taken together, yielded two annotated piRNA candidates. The candidates were both downregulated after ONNV infection as compared to uninfected controls (p=5e-5, q=6.7e-3, locus XLOC_012931, coordinates UNKN:19043685-19043716; and p=9.5e-4, q=0.046, locus XLOC_012762, coordinates UNKN:13088289-13088321; Figure 7).

## Discussion

The current study contributes to a growing body of work defining the deep diversity of the invertebrate virosphere [14, 23, 24]. Because mosquitoes transmit viral infections of humans and animals, there is particular interest in discovery of ISVs comprising the mosquito virome [6, 25-27]. Here, we sampled *Anopheles* mosquitoes from two zones of forest exploitation in Africa and Asia, considered disease emergence zones with likely zoonotic exposure of the human and domestic animal populations. Using assembly quality criteria of non-redundant contigs at least 500 nt in length, we identified 125 novel RNA virus assemblies by sequence similarity to known virus families, and an additional 39 high-confidence virus assemblies that were unclassified by sequence similarity, but display characteristic products of RNAi processing of replication intermediates. Finally, 1566 unclassified contigs possessed comparable assembly quality, and lacked a strong RNAi processing signature, but displayed a signature consistent with piRNA origin. This latter group will require additional work to filter bona fide virus-derived piRNA sequences, which have been previously reported in *Aedes* mosquitoes [17, 18, 20, 21], from other potential sources of piRNAs such as retrotransposons and DNA transposable elements, as well as possible physical degradation.

Nevertheless, taken together at least 164 novel and non-redundant virus assemblies, and possibly many more, were identified in wild *Anopheles* mosquitoes in the current report. Small and long RNAs were sequenced from pools of 5-10 mosquitoes. Pooled sample analysis obscures the distribution and abundance of viruses among individuals in the population. Individual mosquito analysis will likely become a research focus as sequencing costs drop. However, some insight about virus distribution can be gained from comparison of sample pools collected from the same site, for example Senegal or Cambodia. The abundance heat map shown in Figure 5 indicates that virus diversity is high in the population, and evenness is relatively low among sample pools from the same site. This suggests that the number of viruses per individual is probably also low, with a patchy distribution among individuals. This expectation is consistent with a small number of individual mosquitoes with RNAs deep sequenced and de novo assembled in our laboratory, which identifies <5 distinct viruses per individual.

The dynamics of the virome may thus be different from the bacterial microbiome, in which tens of taxa are typically present per individual, and microbial diversity is thought to lead to homeostasis or resilience of the microbiota as an ecosystem within the host [28, 29]. By comparison, very little is known about the function of the mosquito virome within the host. At least three important topics are worth exploring. First, unlike the bacterial microbiota, the stability and resilience over time of the viral assemblage in an individual mosquito is unknown. Members of the virome could persist in individual host populations over time in commensal form, or the uneven and patchy viral distribution observed among sample pools could be a consequence of successive waves of epidemic infection peaks and valleys passing through local populations. The commensal or epidemic models could have distinct biological implications for the potential influence of the virome, including on host immunity and competence for transmission of pathogens.

Second, the individual and population-level effect of ISV carriage on vector competence for pathogen transmission is a key question. In the current study, the predominant host species sampled are *Anopheles* vectors of human malaria, and in Africa, some of these species are also vectors of ONNV. ISVs have not been tested for influence on *Plasmodium* or ONNV infection in *Anopheles*, to our knowledge. ISVs could affect host immunity and malaria susceptibility, or even cause temporary vector population reduction during a putative ISV epidemic. A similar concept may apply to ISV interactions with the mosquito host for arbovirus transmission [26]. We identified relatives of Phasi Charoen-like virus (PCLV) in *Anopheles* from Senegal and Cambodia. PCLV relatives also infect *Aedes*, where they were observed to reduce the replication of ZIKV and DENV arboviruses [30]. Palm Creek virus, an insect specific flavivirus, causes reduced replication of the West Nile virus and Murray Valley encephalitis arboviruses in *Aedes* cells [31]. In any case, ISV co-infection of mosquito vectors with *Plasmodium* and/or arboviruses in nature is highly probable as a general case, because all *Anopheles* sample pools in the current work were ISV-positive, so more research is warranted.

Third, characterization of the arthropod virome may shed light on the evolution of mosquito antiviral immune mechanisms, as well as the evolution of pathogenic arboviruses. ISV replication is restricted to insect cells, but the potential of most mosquito-associated viruses for transmission to humans or other vertebrates is currently unknown, because few studies of host range and transmission have been done. Some viruses may have a host range restricted to only *Anopheles*. For example, Anopheles cypovirus and Anopheles C virus replicate and are maintained by vertical transmission in *An. coluzzii*, but were not able to infect *Ae. aegypti* in exposure experiments [4]. Both of these viruses were able to replicate in *Anopheles stephensi* after exposure, but Anopheles C virus was not stably maintained and disappeared after several generations. Thus, these two viruses may be *Anopheles*-specific, and possibly restricted only to certain *Anopheles* species.

It is likely that the main evolutionary pressure shaping mosquito antiviral mechanisms in general is their persistent exposure in nature to members of the natural virome, rather than the probably less frequent exposure to vertebrate-pathogenic arboviruses. Maintenance of bacterial microbiome commensals in the non-pathogenic commensal state requires active policing by basal host immunity [32]. By analogy, the maintenance of persistent ISVs as non-pathogenic may also result from a dialog with host immunity. Presumably, the same antiviral mechanisms used in basal maintenance of ISVs are also deployed against arboviruses when encountered, which are often in the same families as members of the insect virome [2]. Knowledge of the mechanisms that allow *Anopheles* to carry a natural RNA virome, but apparently reject arboviruses, may provide new tools to raise the barrier to arbovirus transmission by the more efficient *Aedes* and *Culex* vectors.

In addition to the canonical immune signaling pathways, piRNAs can be involved in antiviral protection, although this research is just beginning [18, 33]. One function of genomic piRNA clusters appears to be storage of a molecular archive of genomic threats such as transposable elements, linked to an effector mechanism to inactivate them. This is analogous to bacterial molecular memory mediated by the CRISPR/Cas system. We identified two candidate piRNAs that are downregulated upon ONNV infection in *An. coluzzii*. Involvement of piRNAs during viral infection has not been previously demonstrated in *Anopheles*. piRNA monitoring of the virome may be part of the normal basal management of ISVs, which could potentially be pathogenic if not controlled, but more work is required to draw these connections.

The current report shows that the *Anopheles* virome is complex and diverse, and can be influenced by the geography of mosquito species. This is exemplified by the fact that some viruses are restricted to Senegalese *Anopheles* and others to *Anopheles* from Cambodia (Table3). Similar results were seen in *Ae. aegypti*, where five ISVs were specific to the Australian host population, while six others were found only in the Thai host population [34]. Differences in the *Anopheles* virome across geography could be explained by climate, environmental conditions, breeding sites, and mosquito bloodmeal sources, among other factors. The presence in this study of such a large number of novel and unclassified virus assemblies highlights the fact that the malaria vector virome is understudied. The same observation has been made during metagenomics surveys in *Drosophila*, *Aedes* and *Culex* [24, 35, 36] among other arthropods, indicating that the vast majority of insect viruses are not yet discovered.

## Methods

### Sample collections

Mosquitoes were collected in Cambodia in Kres village, Ratanakiri province (sample pools Cam5−02 and Cam10−02) and Cheav Rov village, Kampong Chnang province (sample pools Cam5−01 and Cam10−01). The majority of inhabitants are engaged in forest-related activities (agriculture, logging and hunting) and may spend the night in forest plots during the harvest period. Vegetation varies from evergreen forest to scattered forest, and the dry season typically runs from November to May and the rainy season from June to October. In Senegal, sampling sites were located in the department of Kedougou in southeastern Senegal. Kedougou lies in a transition zone between dry tropical forest and the savanna belt, and includes the richest and most diverse fauna of Senegal. Recent arbovirus outbreaks include Chikungunya in 2009-2010, Yellow Fever in 2011, Zika in 2010, and Dengue in 2008-2009.

Permission to collect mosquitoes was obtained by Institut Pasteur Cambodia from authorities of Ratanakiri and Kampong Chnang, and by Institut Pasteur Dakar from authorities of Kedougou. Wild mosquitoes visually identified as *Anopheles* spp. at the collection site (non-*Anopheles* were not retained) were immediately transferred into RNAlater stabilization reagent kept at 4°C, and then returned to the laboratory and stored at −80°C until RNA extraction.

### RNA extraction, library construction, and sequencing

Total RNA was extracted from four pools of mosquitoes from each of Senegal and Cambodia (Senegal sample pools: 5 mosquitoes, Dak5-03, Dak5-04, 10 mosquitoes, Dak10-03, Dak10-04; Cambodia sample pools: 5 mosquitoes, Cam5-01, Cam5-02, 10 mosquitoes, Cam10-01, Cam10-02) using the Nucleospin RNA kit (Macherey-Nagel) following the supplied protocol. Library preparation and sequencing steps were performed by Fasteris (Plan-les-Ouates, Switzerland, www.fasteris.com). Long RNA libraries from the eight mosquito pools were made from total RNA depleted of ribosomal RNA by treatment with RiboZero (Illumina, San Diego, CA). Libraries were multiplexed and sequenced on a single lane of the Illumina HiSeq 2500 platform (Illumina, San Diego, CA) by the paired-ends method (2×125 bp), generating on average 36 million high-quality read pairs per library. Small RNA libraries with insert size 18-30 nt were generated from the same eight mosquito pools as above, multiplexed and sequenced in duplicate (two technical replicates per pool) in two lanes of the Illumina HiSeq2500 platform (Illumina, San Diego, CA) by the single-end method (1×50 bp) generating on average 34 million reads of high-quality small RNA reads per library.

### Pre-processing of long and small RNA libraries

Cutadapt 1.13 [37] was used for quality filtering and adaptor trimming of reads from long and small RNA libraries. Low-quality 3’ ends of long RNA reads were trimmed by fixing a phred quality score of 15, and reads smaller than 50 bp after quality filtering and adaptor trimming were removed. In the case of small RNA libraries, reads shorter than 15 bp after quality filtering and adaptor trimming were removed.

In order to filter sequences originating in the mosquito host, sequences passing the above quality filter step were mapped against a custom database consisting of 24 *Anopheles* genomes available in Vectorbase in February 2016 [38]. Bowtie 1.2.0 [39] was used to map small RNA libraries with two mismatches allowed, whereas the BWA-MEM algorithm from BWA-0.7.12 [40] with default parameters was used to map long RNA libraries. Sequence reads that did not map against *Anopheles* genomes, herein referred to as non-host processed reads, were retained and used for de novo assembly and subsequent binning of virus transcripts.

### Estimation of Anopheles species composition of mosquito sample pools

Quality-filtered long RNA read pairs were mapped with SortMeRNA [41] against a custom database of *Anopheles* sequences of the mitochondrial cytochrome c oxidase subunit 1 gene (COI-5P database) extracted from the Barcode of Life database [42]. 98% identity and 98% alignment coverage thresholds were fixed for the operational taxonomic unit (OTU) calling step of SortMeRNA. OTU counts were collapsed at species level and relative abundances of *Anopheles* species with at least 100 reads and 1% frequency in the sample pool were represented as piecharts using the ggplots2 R package.

### De novo sequence assembly and identification of virus contigs by sequence similarity

Processed reads from each country (Cambodia and Senegal) were combined and de novo assembled using different strategies for long and small RNA libraries. Small RNA reads were assembled using the Velvet/Oases pipeline [43] using a range of k-mer values from 13 to 35. Long RNA reads were assembled using both the Velvet/Oases pipeline with a range of k-mer values from 11 to 67 and Trinity [44].

Contigs produced by parallel assembly of Cambodia and Senegal processed reads were filtered in order to remove trans-self chimeric sequences using custom shell scripts, and the resulting contigs were merged with cd-hit-est [45] (95% nucleotide identity over 90% alignment length) in order to generate a final set of non-redundant contig sequences. Non-redundant contigs longer than 500 nucleotides were compared against the GenBank protein sequence reference database using BLASTX [46] with an e-value threshold of 1e-10, and the results were imported into MEGAN6 in order to classify contigs taxonomically using the LCA algorithm [47]. Contigs of viral origin were further manually curated by comparing their sequence with that of the closest virus reference genomes by using Artemis Comparison Tool [48].

### Structural and functional annotation of virus assemblies

Assembled contigs of viral origin were annotated as follows: ORFs were predicted with MetaGeneMark [49], and functionally annotated using Prokka [50] with Virus kingdom as primary core reference database for initial BLASTP searches and including also as reference Hidden Markov Models (HMMS) of virus protein families defined in vFam database [51]. Also, protein sequences of predicted ORFs were processed with the Blast2GO pipeline [52], that generates functional annotation of proteins from BLASTP results against the virus subdivision of GenBank as well as Gene Ontology annotations from top BLASTP results. Prediction of InterPro signatures over viral proteins was also carried out with the InterProScan tool integrated in Blast2GO. The results of the different strategies of structural and functional annotation were integrated and manually curated with Artemis [53].

### Prediction of unclassified contigs of viral origin by small RNA size profiling

In order to recruit contigs of potential viral origin from the pool of unclassified transcripts, we use the approach of Aguiar and collaborators [15]. This approach uses the size profiles of small RNA reads that maps over positive and negative strands of viruses detected by sequence similarity as a signature to identify unclassified transcripts by sequence similarity of potential viral origin. For this purpose, processed small RNA reads were re-mapped over virus contigs and unclassified contigs by sequence similarity using bowtie 1.2.0 [39] allowing at most one mismatch. From the mapped small RNA reads over each contig, the small RNA size profiles were defined as the frequency of each small RNA read of size from 15 to 35 nucleotides that map over the positive and negative strand of the reference sequence. To compute these small RNA size profiles, reads mapped over positive and negative strands of each reference sequence were extracted with Samtools [54], and the size of small RNA reads were computed with the Infoseq program of the EMBOSS package [55]. Custom shell scripts were used to parse Infoseq output to a matrix representing the frequency of reads of different sizes and polarity across virus/unclassified contigs. This matrix was further processed in R (version 3.3.2). In order to normalize the small RNA size profiles, a z-score transformation is applied over the read frequencies of each contig (virus/unclassified). The similarity between small RNA size profiles of virus and unclassified contigs is computed as the Pearson correlation coefficient of the corresponding z-score profiles, and the relationship between small RNA size profiles of virus/unclassified contigs was defined from this similarity values using UPGMA as linkage criterion with the R package Phangorn [56]. These relationships were visualized as heatmaps of the z-score profiles in R with gplots package (version 3.0.1) using the UPGMA dendrogram as the clustering pattern of virus/unclassified sequences. Unclassified contigs with a Pearson correlation coefficient of at least 0.9 with virus contigs and coming from the same mosquito sample pool were regrouped into clusters.

### Phylogenetic analyses

In order to place the new virus sequences characterized in the present study into an evolutionary context, the peptide sequences of RNA dependent RNA polymerase ORFs detected in the annotation step were aligned with the corresponding homologs in reference positive-sense and negative-sense single-strand RNA viruses (ssRNA) and double strand RNA viruses (dsRNA) using MAFFT v7.055b with the E-INS-i algorithm [57]. Independent alignments were generated for all ssRNA and dsRNA viruses and for different virus families (Bunya-Arenavirus, Monenegavirus, Orthomyxovivirus, Flavivirus, Reovirus). The resulting alignments were trimmed with TrimAI [58] in order to remove highly variable positions, keeping the most conserved domains for phylogenetic reconstruction. Phylogenetic trees were reconstructed by maximum likelihood with RAxML [59] with the WAG+GAMMA model of amino acid substitution and 100 bootstrap replicates. Phylogenetic trees were visualized with the R package Ape [60].

### ONNV infection and candidate piRNA gene regulation

Infection of *An. coluzzii* with ONNV, library preparations, and sequencing were described [61]. Briefly, small RNA sequence reads from 2 pools of 12 mosquitoes each fed an ONNV-infected bloodmeal (unfed mosquitoes removed), and 2 control pools of 12 mosquitoes each fed an uninfected normal bloodmeal were mapped to the *An. gambiae* PEST AgamP4 genome assembly using STAR version 2.5 with default parameters [62]. The resulting SAM files were analyzed using featureCounts [63] with default parameters to count mapped small RNAs overlapping with previously annotated *An. coluzzii* piRNA genes in 187 genomic piRNA clusters, in the file, GOL21-bonafide-piRNAs-24-29nt.fastq, from [19]. featureCounts considers a small RNA sequence read as overlapping a piRNA feature if at least one base of the small RNA read overlaps the piRNA feature. Small RNA sequence reads are not counted if they overlap more than one piRNA feature. piRNAs in *An. coluzzii* are annotated by George et al. [19] as novel genes (denoted XLOC loci) as well as piRNAs produced from loci within existing genes of the *An. gambiae* PEST reference (AGAP loci). The Cuffdiff function in Cufflinks version 2.2.1 and DESeq2 version 1.20.0 were used to count and test for significant differential expression levels between ONNV infected and control uninfected samples, yielding 86 piRNA features that were potentially differentially represented in the small RNA sequences between the ONNV and control treatment conditions (Additional File 4: Table S2). The 86 candidates were filtered for a) length of the contiguous region expressed in small RNA less than 40 nt, and b) in the upper 10% of small RNA sequence read depth in all sequence samples combined.

## Declarations

### Ethics approval and consent to participate

There were no human or animal subjects. Permission to collect wild mosquitoes was obtained by Institut Pasteur Cambodia from authorities of Ratanakiri and Kampong Chnang, Cambodia; and by Institut Pasteur Dakar from authorities of Kedougou, Senegal.

### Consent for publication

Not applicable.

### Availability of data and material

All sequence files are available from the EBI European Nucleotide Archive database (http://www.ebi.ac.uk/ena/) under study accession number [REQUESTED], and sample accession numbers: [REQUESTED]). All assembled sequences are available from NCBI (accession numbers: [REQUESTED]).

### Authors’ contributions

Conceived and designed the experiments: EB, FNM, KE, GC, IH, MD, DD, AV, SK, IVS, KDV

Performed the experiments: EB, FNM, KE, GC, IH, MD, DD, AV, SK

Analysed the data: EB, FNM, KE, IVS, KDV

Wrote the manuscript: EB, FNM, KE, IVS, KDV

All authors read and approved the final manuscript.

## Acknowledgements

We acknowledge the assistance of Allan Dickerman, Jiyoung Lee and Song Li, Virginia Polytechnic Institute and State University, and of Silke Jensen, Clermont Université, Clermont-Ferrand, France, for helpful advice and assistance on piRNA analysis.

## Supporting information

**Additional File 1: Table S1.**
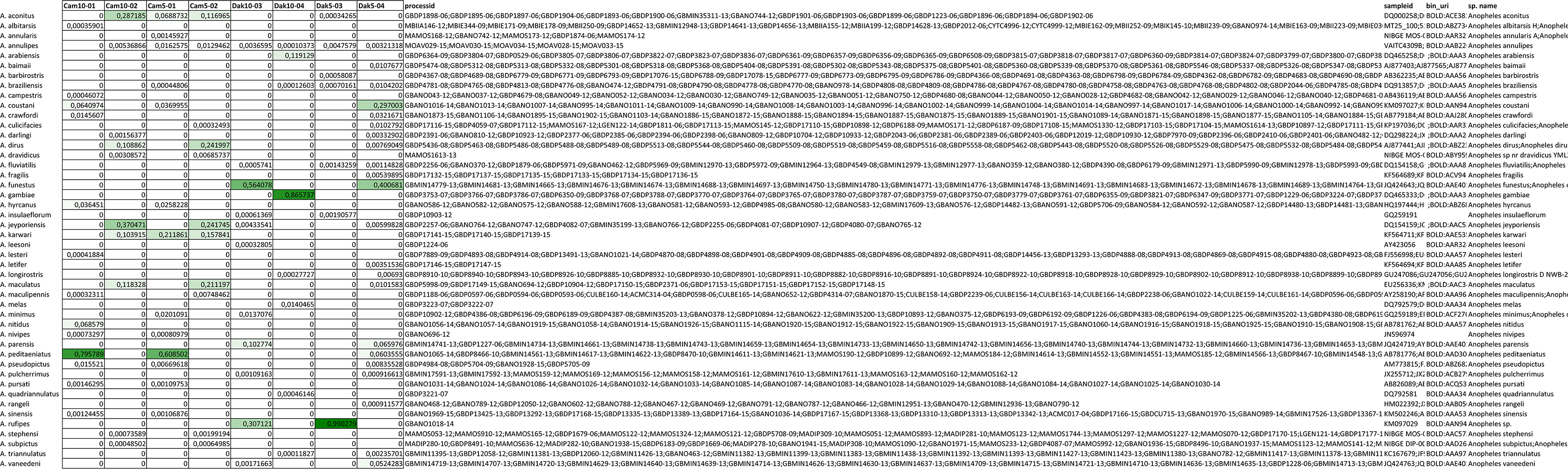
*Anopheles* mosquito taxa represented in the collections from Senegal and Cambodia, as detected by comparison to *Anopheles* sequences from the Barcode of Life COI-5P database. Data corresponds to pie charts of *Anopheles* taxa by country and sample pool depicted in Figure 1.

**Additional File 2: Figure S1.**
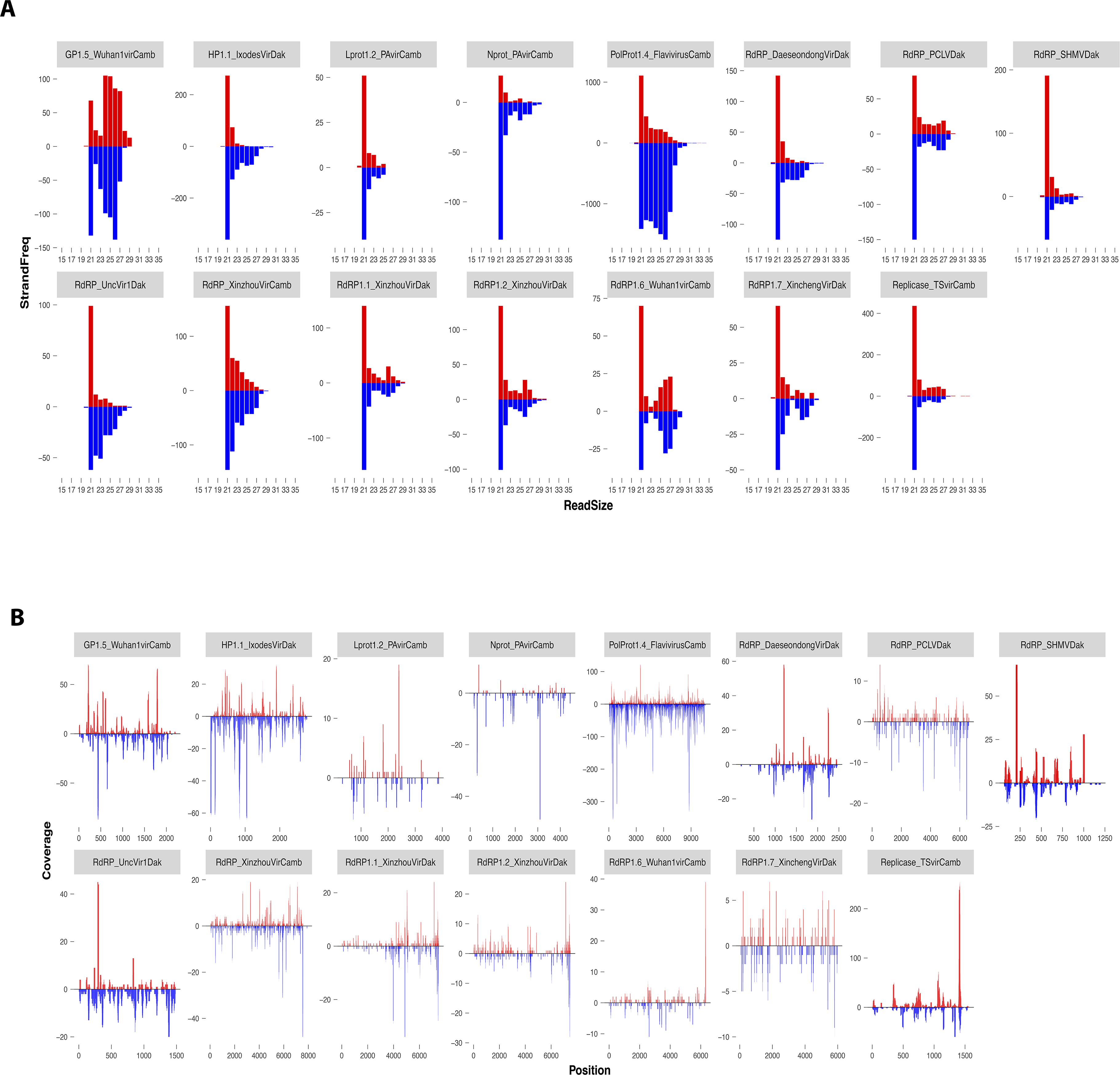
Small RNA size profiles (A) and coverage profiles (B) of 15 novel virus assemblies with classic RNAi processing pattern. Virus assemblies shown are in Figure 5, Cluster 3, and are classified by sequence similarity to known virus assemblies. Red vertical bars represent reads mapped over the positive strand of reference viral sequence, and blue bars represent reads mapped over the negative strand.

**Additional File 3: Figure S2.**
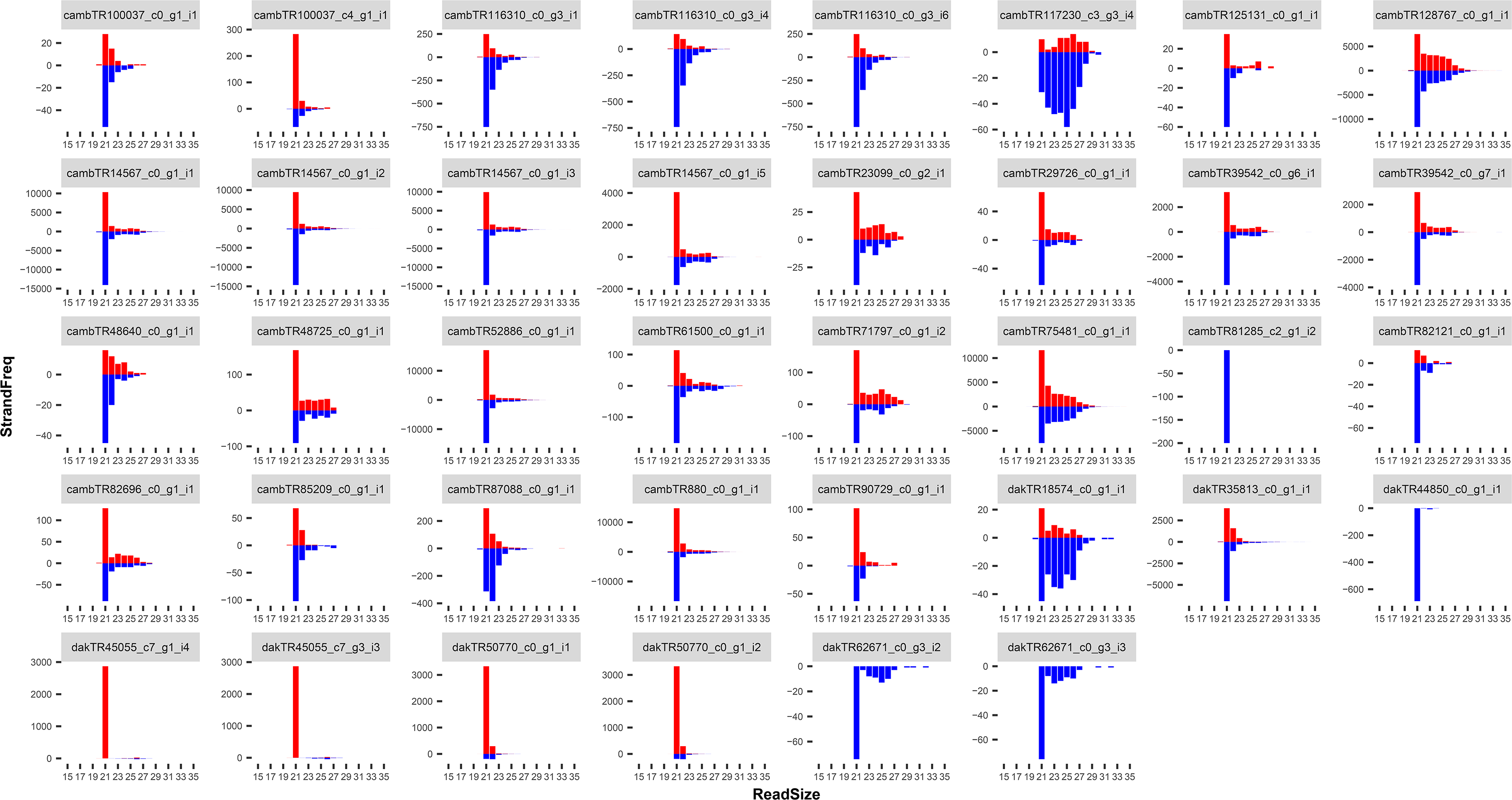
Small RNA size profiles of contigs left unclassified by sequence similarity grouping. Unclassified contigs that display strong association by small RNA profile with Figure 5, Cluster 2 and Cluster 3. Red bars represent reads mapped over the positive strand of reference viral sequence, and blue bars represent reads mapped over the negative strand.

**Additional File 4: Table S2.**
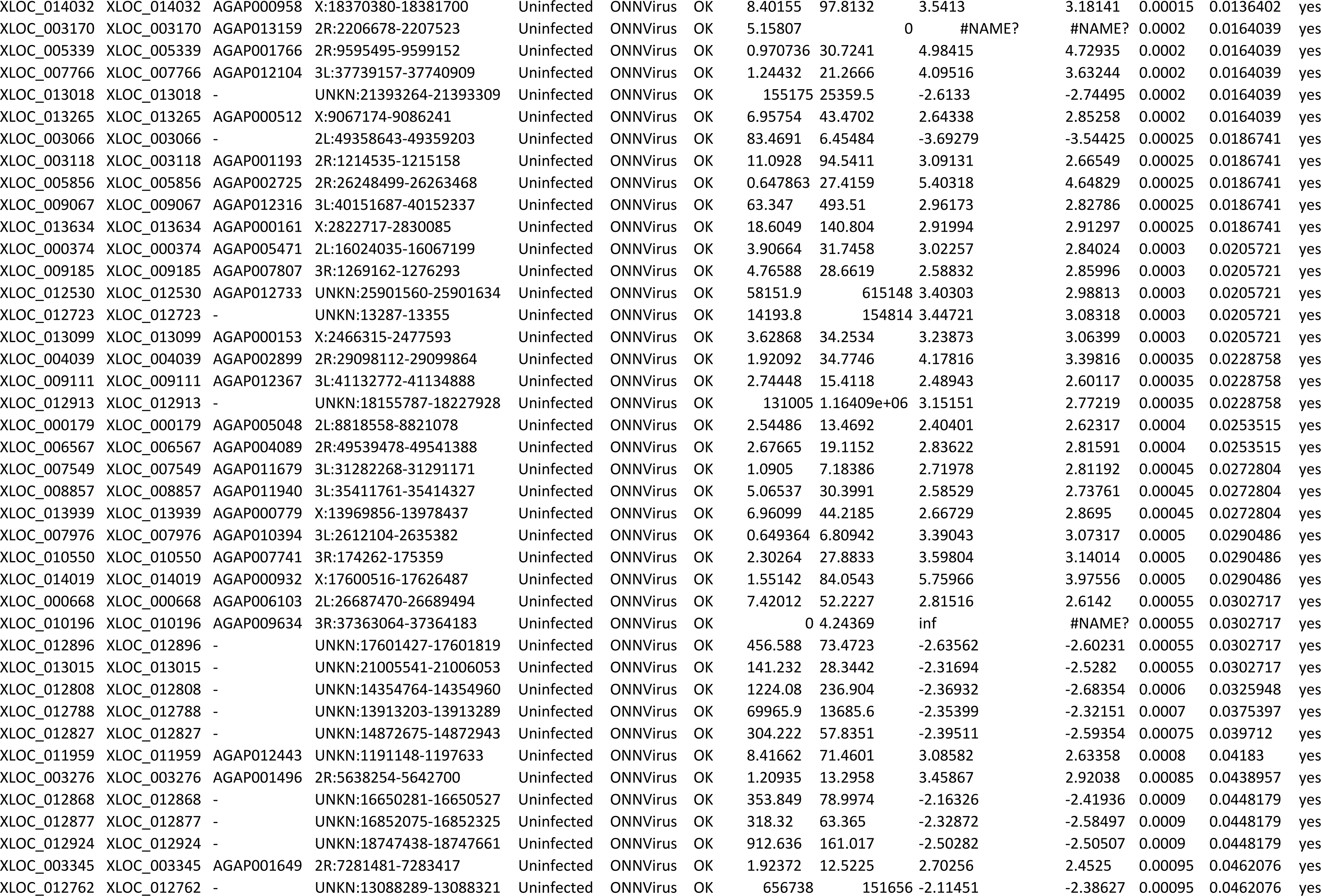
Anopheles coluzzii piRNAs potentially differentially represented in the small RNA sequences between the ONNV and control treatment conditions.

